# Anti-apoptotic clone 11 derived peptides induce in vitro death of CD4+ T cells susceptible to HIV-1 infection

**DOI:** 10.1101/2020.04.06.028860

**Authors:** Anastassia Mikhailova, José Carlos Valle-Casuso, Annie David, Valérie Monceaux, Stevenn Volant, Caroline Passaes, Amal Elfidha, Michaela Müller-Trutwin, Jean-Luc Poyet, Asier Sáez-Cirión

**Affiliations:** Institut Pasteur, Unité HIV Inflammation et Persistance, Paris, France; Institut Pasteur, Hub Bioinformatique et Biostatistique – C3BI, USR 3756 IP CNRS – Paris, France; INSERM UMRS1160, Institut Universitaire d’Hématologie, Hôpital Saint-Louis, Paris, France; Université Paris Diderot, Université de Paris, Paris, France; Université Paris Descartes, Université de Paris, Paris, France

## Abstract

HIV-1 successfully establishes long-term infection in its target cells despite viral cytotoxic effects. We have recently shown that cell metabolism is an important factor driving CD4+ T-cell susceptibility to HIV-1 and the survival of infected cells. We show here that expression of anti-apoptotic clone 11 (AAC-11), an anti-apoptotic factor upregulated in many cancers, increased with progressive CD4+ T cell memory differentiation in association with the expression of cell cycle, activation and metabolism genes and correlated with susceptibility to HIV-1 infection. Synthetic peptides based on the LZ domain sequence of AAC-11, responsible for its interaction with molecular partners, were previously shown to be cytotoxic to cancer cells. Here we observed that these peptides also blocked HIV-1 infection by inducing cell death of HIV-1 susceptible primary CD4+ T-cells across all T-cell subsets. The peptides targeted metabolically active cells and had the greatest effect on effector and transitional CD4+ T cell memory subsets. Our results suggest that AAC-11 survival pathway is potentially involved in the survival of HIV-1 infectable cells and provide a proof of principle that some cellular characteristics can be targeted to eliminate the cells offering the best conditions to sustain HIV-1 replication.

**IMPORTANCE:** Although antiretroviral treatment efficiently blocks HIV multiplication, it cannot eliminate the cells already carrying integrated proviruses. In the search for a HIV cure the identification of new potential targets to selectively eliminate infected cells is of the outmost importance. We show here that peptides derived from the anti-apoptotic clone 11 (AAC-11), which expression levels correlated with susceptibility to HIV-1 infection of CD4+ T-cells, induced cytotoxicity in CD4+ T-cells showing the highest levels of activation and metabolic activity, conditions known to favor HIV-1 infection. Accordingly, CD4+ T-cells that survived the cytotoxic action of the AAC-11 peptides were resistant to HIV-1 replication. Our results identify a new potential molecular pathway to target HIV-1 infection.

## INTRODUCTION

Human immunodeficiency virus type 1 (HIV-1) is a persistent viral infection that has claimed millions of lives around the globe. The era of combination anti-retroviral therapy (cART) has extended life expectancy of people living with HIV and improved their quality of life. However, although cART blocks viral replication, it does not eliminate infected cells, which persist despite decades of treatment and originate viral rebound if therapy is interrupted (1–3). There is, therefore, a need to identify characteristics of HIV-infected cells that could be potentially targeted by new therapeutic strategies. CD4+ T cells are primary targets of HIV-1, however these cells differ in their relative susceptibility to infection (4–6). HIV requires a particular cell environment providing abundant factors that the virus exploits to sustain its replication cycle. Susceptibility to HIV infection *in vitro* increases with CD4+ T cell differentiation. Naïve CD4+ T cells are most resistant while central, transitional and effector memory CD4+ T cells are progressively more susceptible to the virus. We have recently shown that these differences are, at least in part, related to the increased metabolic activity associated with progressive differentiation of these subsets (7). Immunometabolism is a critical element in the regulation of T cell differentiation, survival and function (8). Upon antigenic stimulation T cells upregulate metabolic fluxes to provide the energy necessary to support cellular processes and to increase the pool of substrates necessary for building proteins, lipids, nucleic acids and carbohydrates. This metabolically rich environment is necessary for the establishment of both productive and latent HIV infection (7, 9, 10), as it is also the case for other infections (11–13), and may offer new opportunities to tackle HIV.

In a post-hoc analysis of results obtained in our previous study (7) we found that the anti-apoptotic clone 11 (AAC-11) [also known as apoptosis inhibitor 5, API5] was significantly correlated with infection in different subsets of memory CD4+ T cells. The anti-apoptotic activity of AAC-11 might contribute to the survival of metabolically active cells. Indeed, AAC-11 is overexpressed in many cancers (14) and allows cancer cell survival in conditions of metabolic stress (15). Its expression is associated with poor prognosis in non-small lung and cervical cancers (16–18). Although the mechanisms associated with its anti-apoptotic activity have not been clearly elucidated, AAC-11 contains several protein interaction domains, including a leucine zipper (LZ) domain (19), and has been proposed to repress apoptotic effectors such as E2F1 (20), Acinus (21) and caspase-2 (22). Synthetic peptides based on the LZ domain sequence of AAC-11 were previously shown to be cytotoxic to cancer cells both *in vitro* and in a *in vivo* mouse model of melanoma (23, 24) or acute leukemia (25).

We explored here whether AAC-11 derived peptides could, similarly to its action against cancer cells, induce the elimination of HIV-1 infected cells. We found that AAC-11-derived peptides were preferentially cytotoxic for CD4+ T cells targeted by HIV-1. In contrast, cells escaping the cytotoxic action of the peptides were resistant to HIV-1 replication. These results offer proof of principle that some characteristics of the cells targeted by the virus could be antagonized to counteract infection.

## RESULTS

### AAC-11 derived peptides display anti-viral activity against diverse HIV-1 viral strains and SIV

We previously analyzed the susceptibility of memory CD4+ T cell subpopulations (central (Cm), transitional (Tm) and effector (Em) memory) to HIV-1 infection (7). We also analyzed the expression, at the time of infection, of a panel of 96 genes related to T cell differentiation, function and survival as well as restriction and HIV facilitating factors (https://data.mendeley.com/datasets/vfj3r27gnf/1). Among other genes, we found in post-hoc analysis, the expression of anti-apoptotic factor AAC-11 to increase progressively with the stage of differentiation of memory CD4+ T cells (Cm<Tm<Em) and to correlate with infection in these subsets (Fig 1A,B). The expression of AAC-11 was strongly correlated with the expression of multiple genes related to cell metabolism (e.g. SLC1A5, MTOR, HIF1A, GUSB, GAPDH), cell cycle (e.g. RHOA, RB1, MAPK1, MKI67) and genes known to be necessary for different steps of the HIV replication cycle (RRM2, SAMRCB1, DDB1, CFL1, ACTB, CDK9, NFKB1) (See Table S1 for references to gene functions and/or association with HIV replication) (Fig 1C). These results suggest that AAC-11 anti-apoptotic pathway is upregulated in the cells that offer an optimal environment for HIV-1 replication.

**Figure 1.**
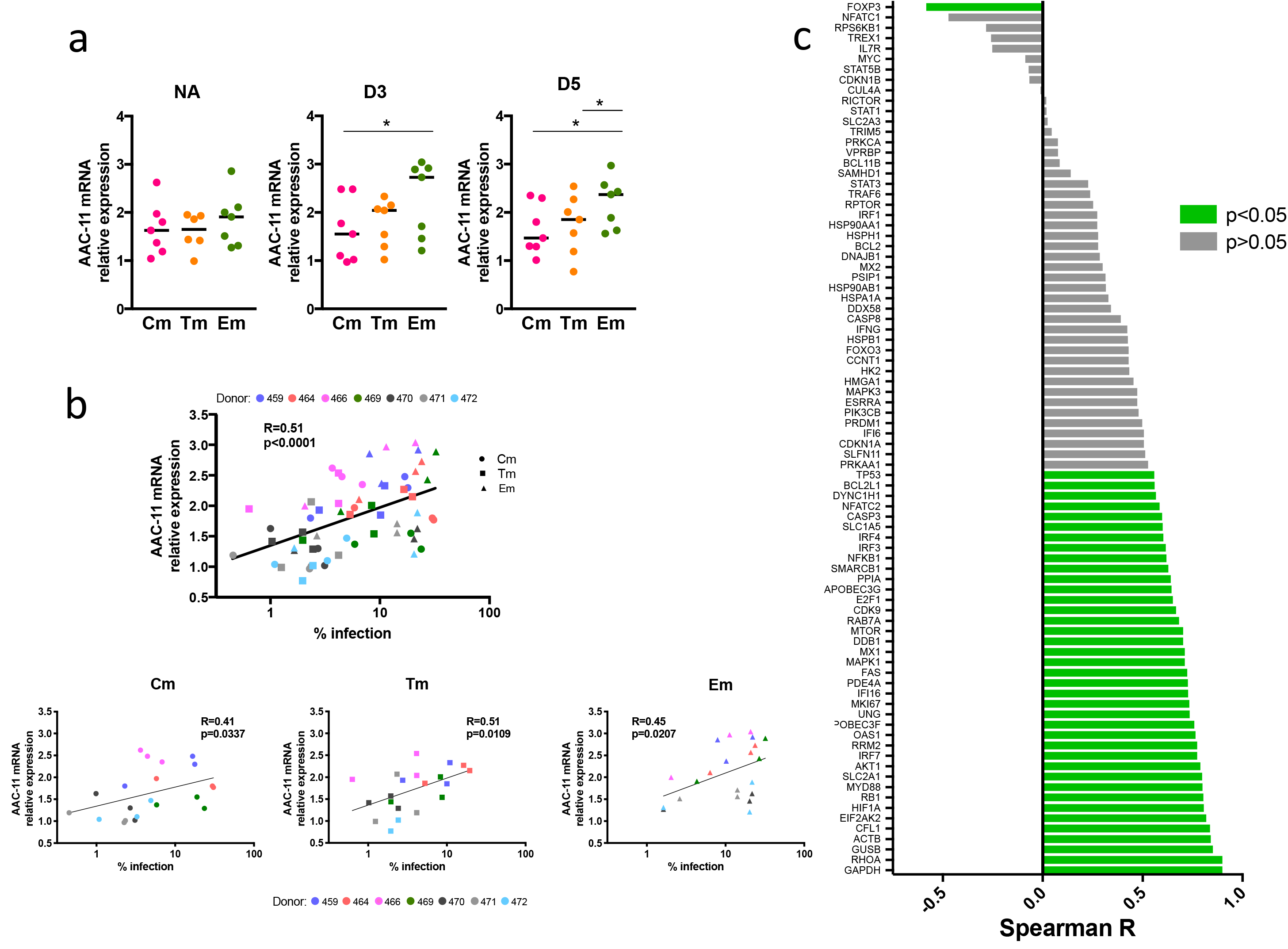
AAC-11 gene expression correlates with the proportion of infection in each memory CD4+ T cell subset and is associated with expression of various cell cycle and metabolism genes. **A.** Non-activated or activated (day 3 and 5) CD4+ T cell memory subsets (Cm, Tm, Em) were analyzed for the expression of AAC-11 at the time of infection. **B.** Non-activated or activated (day 3 and 5) Cm, Tm and Em subsets were analyzed for the levels of AAC-11 gene expression and correlated with % of infected cells in each subset at day 3 post-infection **C.** The correlation of AAC-11 gene expression with other genes in a 96-gene array.

We sought to antagonize the action of AAC-11 by using a panel of synthetic peptides derived from the LZ domain located in its alpha helix 18 (19), fused to a cell penetrating sequence of antennapedia protein of *Drosophila melanogaster* [also commonly known as penetratin] (RK16 here) (Fig 2A), which can be used as an intracellular delivery method for its cargo (26). These AAC-11 derived peptides act as competitive inhibitors of protein-protein interactions, can prevent AAC-11 anti-apoptotic activities and have shown anti-tumor activities *in vitro* and *in vivo* (23–25). RT53 spanned the entire length of the AAC-11 LZ domain (Figure 2A), other peptide spanned progressively smaller region of the LZ domain. We applied these peptides to activated primary human CD4+ T cells (27) immediately after infection with HIV-1 and measured infection levels 3 days later (Fig 2B). AAC-11-derived peptides reduced HIV-1 infection in a concentration-dependent manner (Fig 2C). Shorter peptides were less effective at inhibiting infection (Fig 2C, D). RK16, the penetratin alone, did not show any anti-viral effect even at the highest concentration tested. We observed a rapid drop in the proportion of infected cells at 6μM peptide concentration for RT53 and RT39, peptides exhibiting the strongest anti-viral effect. At this concentration, RT53 was most potent at suppressing infection (EC50=4.19μM) (Fig 2D and Fig S1), indicating that the entire length of the LZ domain of AAC-11 was necessary for full peptide potency. We, therefore, proceeded to work with RT53 at 6μM concentration.

**Figure 2.**
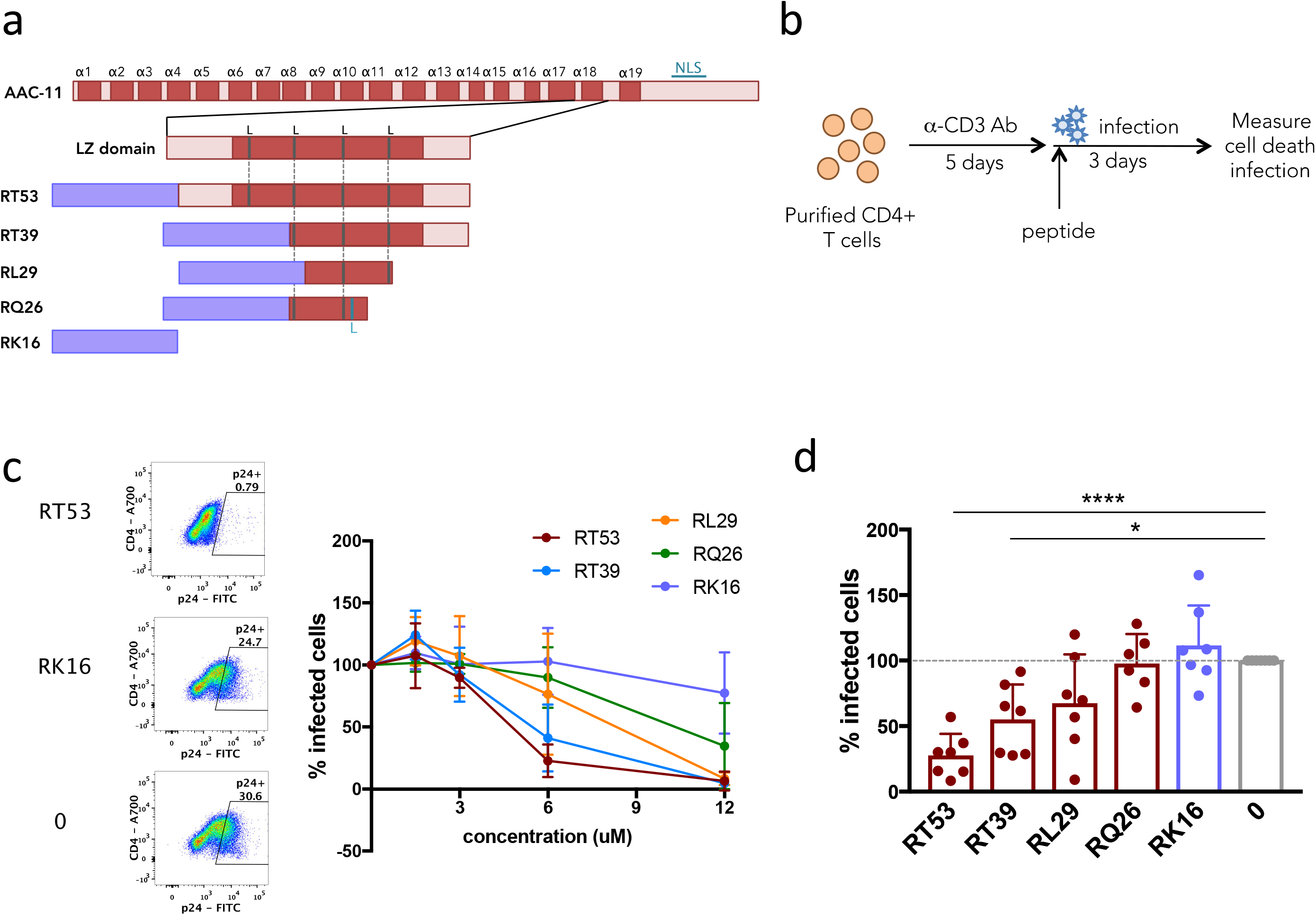
AAC-11-derived peptides inhibit HIV-1 infection. **A.** Schematic representation of AAC-11-derived peptides indicating LZ domain **B.** Experimental scheme of treatment of infected cells with AAC-11-derived peptides **C.** Representative flow cytometry plot of p24 staining in live, untreated or cell treated with 6 μM RT53 or RK16. A dose-response curve of the effect of AAC-11-derived peptides on HIV-1 infection in living cells **D.** The proportion of infection as compared to non-treated control among living cells infected with HIV-1 BaL and treated with 6 μM of indicated peptides.

We next tested if RT53 was able to inhibit infection of distinct HIV-1 viruses (Fig 3A). Both R5 (Bal) and X4 (NL4.3) strains as well as primary isolates (BX08, DH12, 132w) and single cycle VSVG-pseudotyped HIV-1 particles were inhibited, albeit less efficiently for the latter. RT53 was also able to inhibit *in vitro* infection of human CD4+ T cells with simian immunodeficiency virus SIV_mac251_. Moreover, it inhibited viral spread from splenic CD4+ T cells of Cynomologus macaques previously infected with SIV_mac251_ (Fig 3A). The inhibition of infection in both human and macaque cells is coherent with the high conservation of AAC-11 across species (20) and 100% homology between humans and macaques. Again, RK16 alone had no inhibitory effect for any of the viruses tested. Overall, our results show that AAC-11-derived peptides were able to impair HIV-1 and SIV infection of CD4+ T cells *in vitro* as well as spread from *in vivo* infected CD4+ T cells.

**Figure 3.**
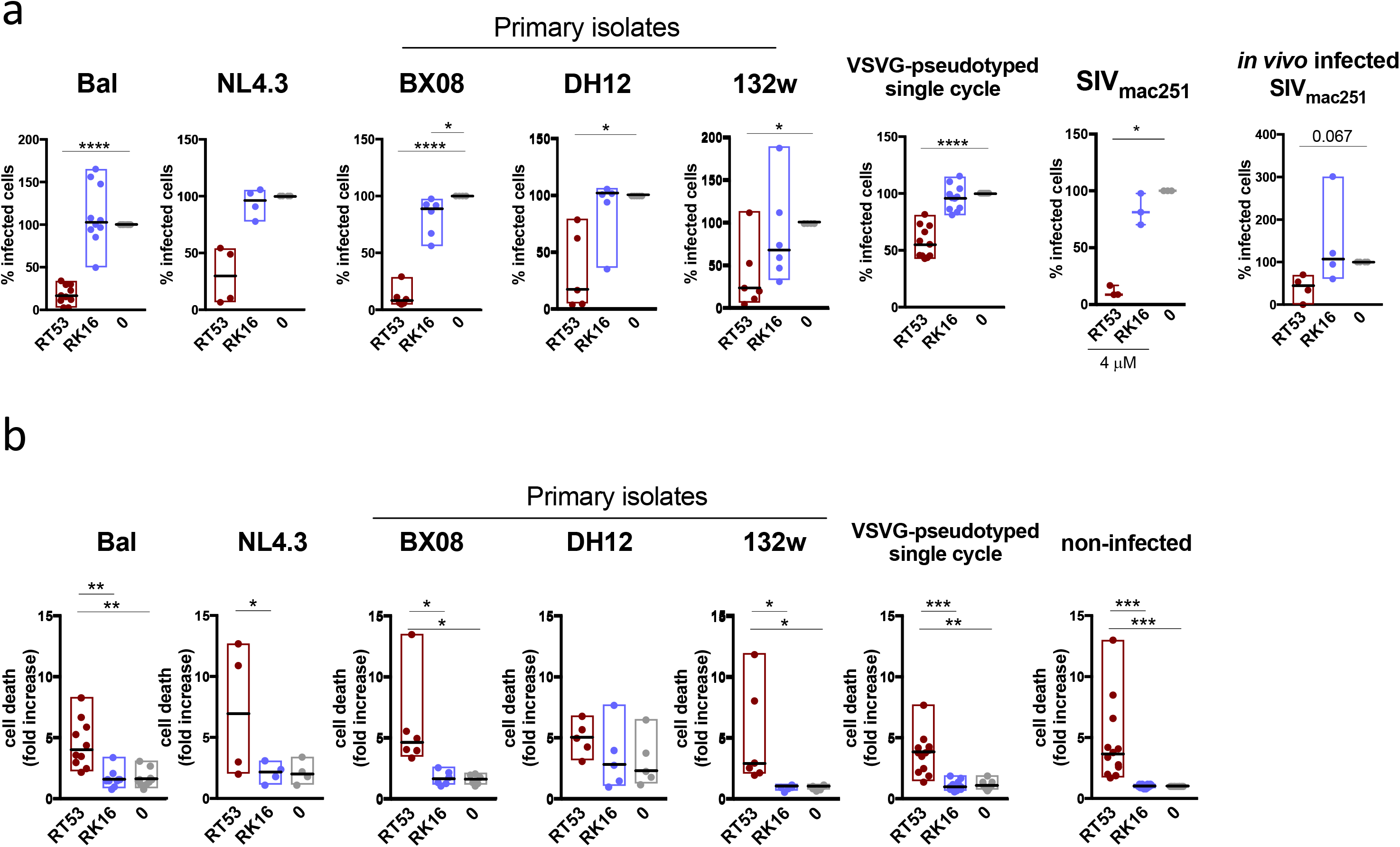
RT53 inhibits infection of diverse HIV-1 strains and SIV. Activated primary CD4+ T cells were infected with different HIV-1 viral strains and incubated with 6 μM RT53 or RK16 for 3 days **A.** Infection was measured by flow cytometry (intracellular p24) or ELISA (p24/p27) in supernatants **B.** Cell death was measured by flow cytometry (LIVE/DEAD™ Violet Viability dye). Values are expressed as a range (min-max) with median indicated. Each symbol represents experiments with cells from a different donor.

### Blockage of HIV-1 infection by AAC-11-derived peptides is associated with cell death

Since the AAC-11-peptides have been shown to induce cytotoxicity in cancer cells (21), we next evaluated the amount of cell death in our cultures. Indeed, we saw that all peptides caused various rates of cell death (Fig 4A). Moreover, we found that the proportion of cell death caused by different peptides in different donors was negatively correlated with the proportion of HIV infected cells in the cultures (Fig 4B). As before, RT53 was most potent at causing cell death. Interestingly, RT53 induced similar death rates in cells infected with various HIV-1 strains and in cells that had not been challenged with HIV-1 (Fig 3B and Fig 4A). This indicated that some CD4+ T cells had pre-existing sensitivity to the action of the peptides that was not dependent on infection.

**Figure 4.**
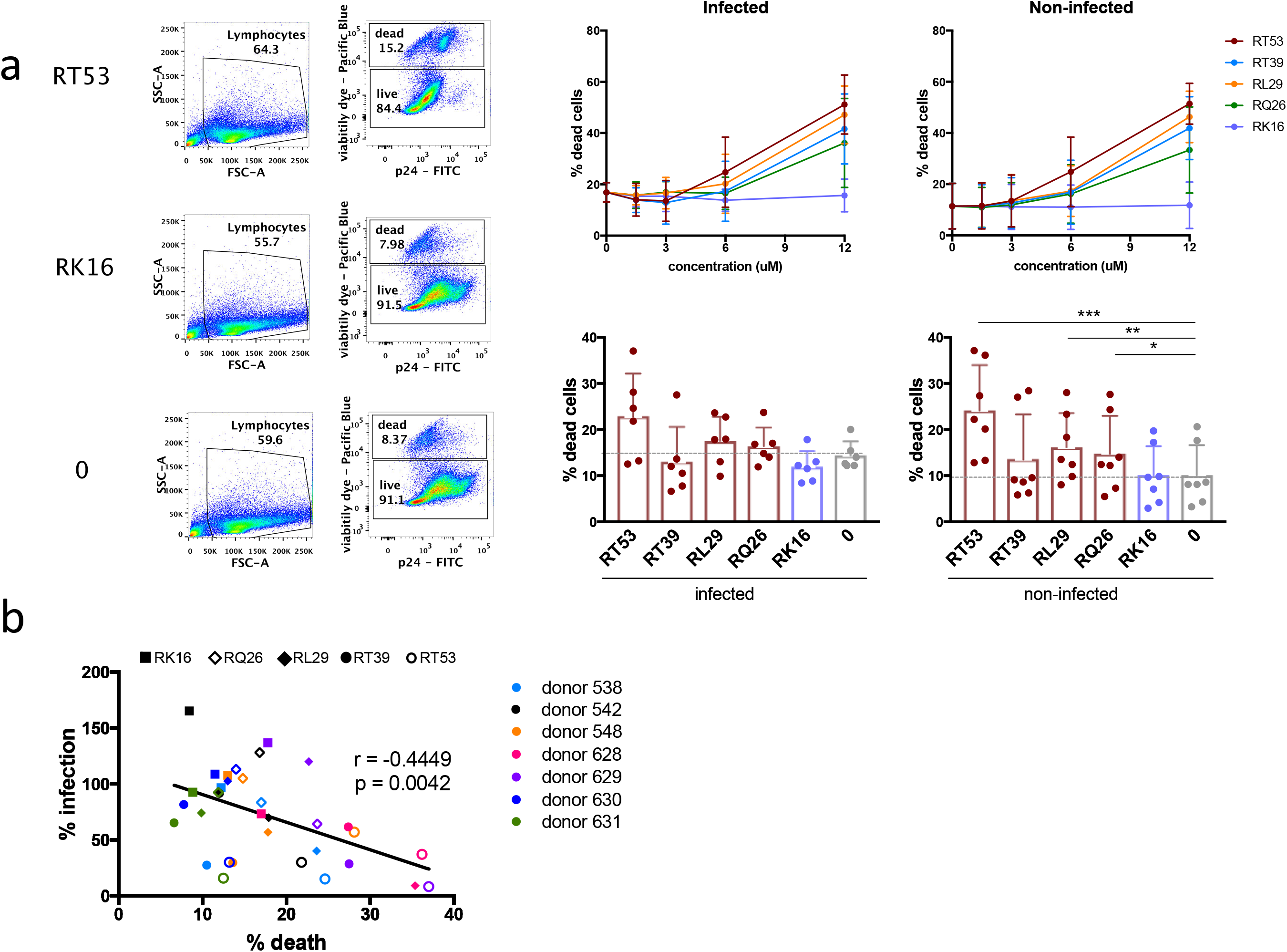
Decrease in HIV-1 infection is associated with cell death. **A**. Example of flow cytometry gating strategy to evaluate cell death in cultures. *In vitro* activated primary CD4+ T cells were incubated with various concentrations of AAC-11-derived peptides after infection with HIV-1 Bal and cell death was evaluated by flow cytometry on day 3 of infection **B**. Proportion of infected cells among live cells vs cell death in different donors’ CD4+ T cells incubated with 6 μM indicated peptides. Each symbol represents experiments with cells from a different donor.

To explore the specificity of action of AAC-11 derived peptides, we analyzed if RT53 induced cell death in other cell types. We found out that CD8+ T-cells were susceptible to the action of RT53, although at lower extent than CD4+ T-cells (p=0.039), while NK cells were resistant to the cytotoxicity of RT53 (p=0.0078) (Fig S2A). Therefore, RT53 did not have a general cytotoxic effect on all lymphocytes. On the other hand, RT53 induced cytotoxicity in highly susceptible Jurkat and SupT1 human CD4+ T cell lines, but it did not affect the proportion of infected cells (Fig S2B), not recapitulating the selective effect observed in primary CD4+ T-cells, further suggesting that the link between the peptide-induced cytotoxicity and antiviral activity is linked to particular properties of primary CD4+ T-cells that favor HIV replication.

Together, these results show that AAC-11 did not exert a non-specific cytotoxic action but affected particular cell subsets. Primary CD4+ T cells preferentially targeted by HIV-1 for infection had an enhanced sensitivity to the action of AAC-11-derived peptides. Hence, AAC-11-derived peptides may induce selective elimination of HIV-1 target cells.

### Cell death induced by AAC-11-derived peptides is associated with K^+^ efflux and mitochondrial depolarization

We next investigated the mechanism of RT53-mediated effect. We started by evaluating the kinetics of RT53 action. We used “real time” flow cytometry to read out changes in several molecular indicators associated with cell death as a function of time (28). We found that the treatment of cells with RT53 induced a sharp and rapid increase in the surface levels of phosphatidylserine (PS) (as determined by binding to annexin V-FITC), classically associated with apoptosis induction (Fig 5A). Additionally, in a subset of RT53-treated CD4+ T cells, we also observed a decrease in the intracellular levels of K^+^ ion (revealed by fluorescent K^+^ indicator 2-APG), the efflux of which has been linked to the activation of the cell death machinery (29–31); and mitochondrial depolarization, evaluated by the increase in green fluorescence of mitochondrial membrane potential dye JC-1 (Fig 5A). These events were accompanied by the increase in DNA labelling with 7-Aminoactinomycin D (7-AAD), a classical marker for dead cells. Cell death peaked at 30 minutes after treatment with the RT53 peptide in both non-infected and infected cells, with no significant increase at later time points (Fig 5B).

**Figure 5.**
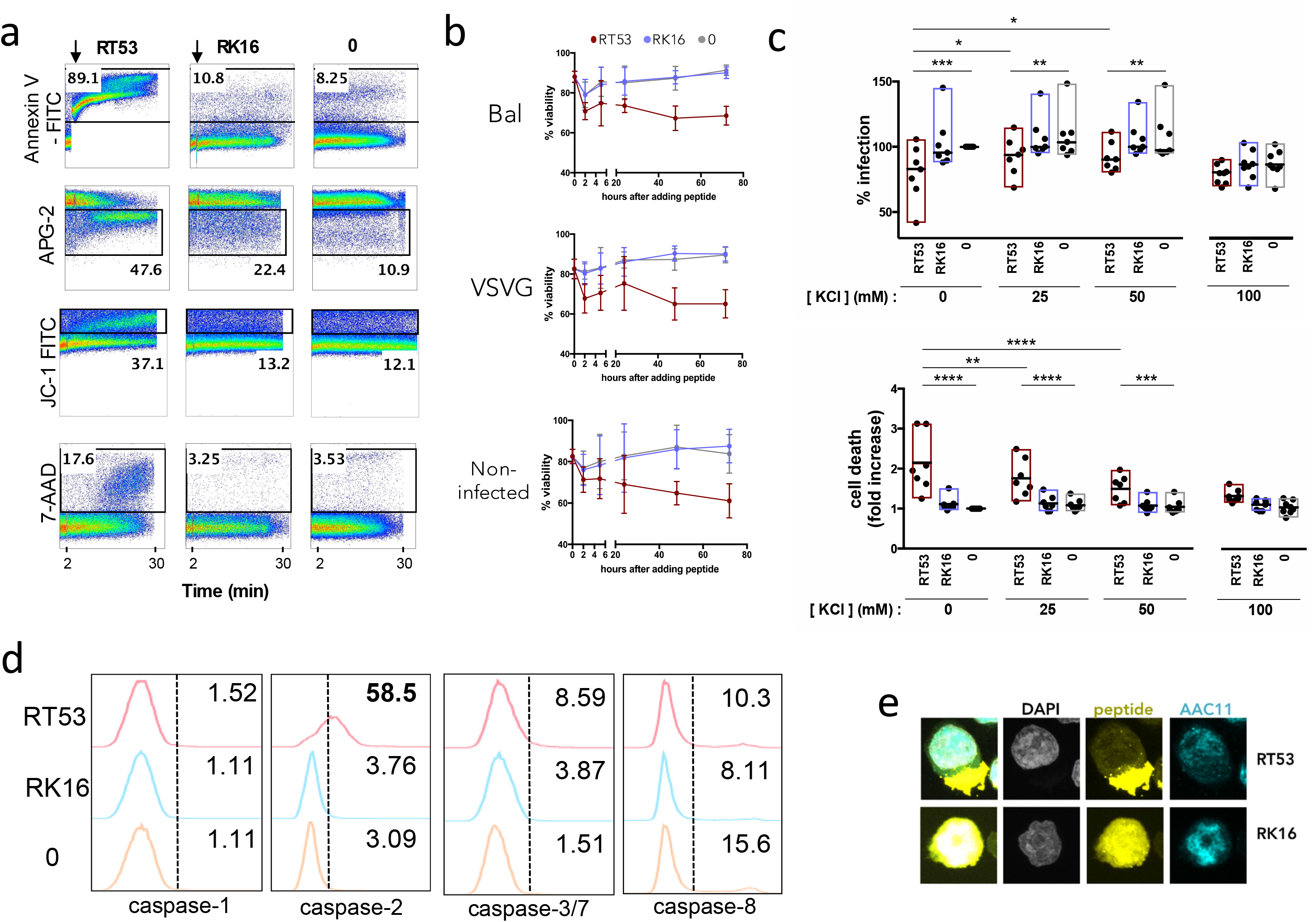
RT53-mediated cell death is rapid and is associated with K^+^ efflux and mitochondrial depolarization. **A.** Cells were placed in Annexin V-FITC and 7-AAD staining buffer or loaded with fluorescent K^+^ indicator (APG-2) or mitochondrial membrane potential probe (JC-1) and run on a flow cytometer to acquire fluorescence over time. Time point of peptide addition (after 2 minutes of acquisition) is indicated with an arrow **B**. Infected cells were incubated with 6 μM RT53 or RK16 and viability determined by trypan blue at indicated time points **C.** Cells pre-infected with VSVG-pseudotyped HIV-1 particles (day 3 of infection) were incubated with 6 μM RT53 or RK16 in the presence of various concentrations of KCl for 5 hours. Infection and cell death were then evaluated by flow cytometry. Symbols represent experiments with cells from different donors. Bars represent min-max range with medians indicated. **D.** Cells were incubated with 6 μM RT53 or RK16 for 2 hours and stained with probes for active caspases **E.** Immunofluorescent detection of RT53/RK16-Rd and AAC-11 in CD4+ T cells. Cells were incubated with 6 μM RT53/RK16-Rd for 2 hours, washed, immobilized on a polylysine coated coverslips, fixed and stained with antibodies for AAC-11 and DAPI.

To investigate if cell death caused by the peptides was indeed responsible for the observed anti-viral effect, we sought to revert RT53-induced cytotoxicity by antagonizing intracellular K^+^ efflux with increasing KCl concentrations in the culture medium (32). Non-infected CD4+ T cells or cells that had been pre-infected with VSVG-pseudotyped HIV particles were incubated with RT53 or RK16 for 5 hours in presence of increasing extracellular K^+^ concentration of up to 100 mM KCl supplemented in the cell culture medium (Fig 5C). We observed a KCl concentration-dependent decrease in RT53-mediated cell death, relative to RK16 or untreated condition, which was accompanied by a progressive increase in the proportion of infected CD4+ T cells recovered at the end of the culture. No differences were observed in the rates of cell death and the percentage of infected cells between RT53 and RK16 or control when the cultures were done in the presence of 100 mM of KCl, although we should notice that at this concentration the rates of infection were overall lower than in CD4+ T cells cultured in the absence of extra KCl (Fig 5C).

We then evaluated if RT53 treatment had an impact on the activation of caspases, known activators and executioners of cell death. Of note, caspase-2 has been recently reported as being repressed by AAC-11 (22). Upon treatment with the AAC-11 derived peptide, we did not observe significant activation of caspase-1, caspase-3/7 and caspase-8, but we saw a strong activation of caspase-2 (Fig 5D). Nevertheless, inhibition of caspase-2 activation did not lead to a decrease in cell death (Fig S3A), suggesting that the activation of caspase-2 was not the main mechanism responsible for cell death in our system. Similar results were obtained with inhibition of caspase-1, caspase-3/7 and caspase-8 (Figure S3B). These results are in agreement with caspase-independent mechanism of cytotoxicity induced by RT53 in cancer cells (24).

RT53 was previously shown to mediate its cytotoxic effect through a membranolytic action by way of the interaction with unknown AAC-11 partner (23, 24). We therefore determined the cellular localization of RT53 and RK16 conjugated to Rhodamine (Rd). As expected RK16-Rd was found in the cytoplasm of the CD4+ T cell (Figure 5E) while RT53-Rd localized predominantly to the cell membrane. These studies were carried out at 6μM peptide concentration but similar results were found at subcytotoxic peptide concentrations of 2 and 4μM (data not shown). Therefore, our results indicate that similar to cancer cells, RT53 could be causing cell death in primary CD4+ T cells by membranolytic action.

Overall, our results show that RT53 induced rapid cell death in a subset of CD4+ T cells possibly through a membranolytic mechanism. The impairment of peptide-induced cell death abrogated the anti-viral activity of the peptides, which provides a direct link between the peptide cytotoxic effect and its capacity to inhibit HIV infection.

### RT53 preferentially depletes effector and transitional memory CD4+ T cell subpopulations

Our previous results showed that AAC-11 mRNA expression was upregulated with CD4+ T cell differentiation (Fig 1). We have now found that the expression of AAC-11 at the protein level also increased with T cell differentiation (Fig 6A, Fig S4). We thus wondered if RT53 might have a distinct impact on different CD4+ T cell subsets. RT53 treatment changed the relative contribution of CD4+ T cell subsets to the total pool of CD4+ T cells when compared to the non-treated or RK16-treated conditions (Fig 6B). Naïve CD4+ T cells were enriched upon treatment with RT53 while the more differentiated Tm and Em CD4+ T cells were depleted. Although activation *in vitro* changed somewhat the relative contribution of CD4+ T cell subsets to the pool of CD4+ T cells, RT53 had a similar impact on CD4+ T cell subsets when we used non-activated or *in vitro* activated cells (Fig 6B). As previously reported (7), in our experimental conditions naïve CD4+ T cells were highly resistant to infection with both VSVG pseudotyped HIV-1 particles and HIV-1 Bal (Fig S5). Susceptibility to infection increased with T cell differentiation, with Em and Tm cells showing the highest infection levels (Fig S5). Thus, our results showed that RT53 preferentially targeted differentiated memory cells which had heightened susceptibility to HIV infection.

**Figure 6.**
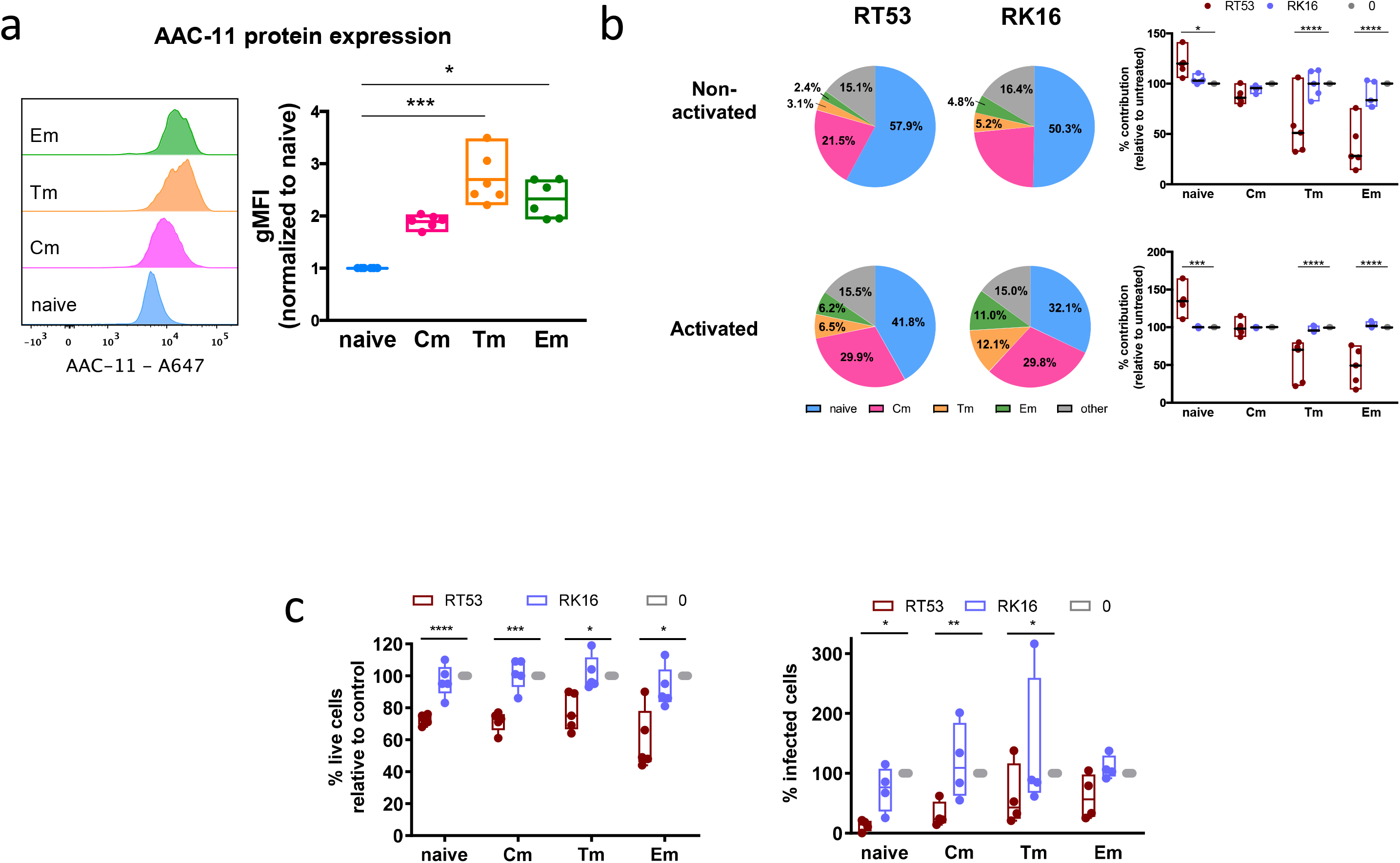
RT53 preferentially targets Tm and Em cells to cell death. **A.** *In vitro* activated CD4+ T cells were stained for markers of CD4+ T cell memory subsets and AAC-11. **B.** *In vitro* activated or non-activated CD4+ T cells were pulsed with 6 μM RT53 or RK16 for 5 hours, harvested and stained for markers of CD4+ T cell memory subsets. **C.** *In vitro* activated CD4+ T cells were sorted using fluorescently activated cell sorting (FACS) into naïve, Cm, Tm and Em subsets. Each subset was pulsed with 6 μM RT53 or RK16 for 24 hours and viability was measured by flow cytometry with LIVE/DEAD™ Violet Viability dye. In parallel, sorted cells were infected with HIV-1 Bal in the presence or absence of HIV-1 Bal and rates of infection were determined on day 3 post-infection by intracellular p24. Viability and infection levels are expressed in relation to the untreated condition for each individual.

However, not all Em and Tm cells were depleted after RT53 treatment and some extent of peptide-induced cell death could also be observed in other CD4+ T cell subsets, including naïve cells. We therefore separately tested the impact of RT53 treatment on isolated naïve, Cm, Tm and Em CD4+ T cells. We confirmed that RT53 induced cell death in each of the CD4+ T-cell subsets (Fig 6C). Moreover, RT53 treatment decreased HIV-1 infection in all CD4+ T-cell subsets. Overall, these results show that RT53 targeted cells with specific characteristics that were strongly enriched, but not exclusively found, within more differentiated subsets. Of note, the cells targeted by RT53 within each subset were susceptible to HIV-1 infection.

### RT53 targets metabolically active cells

We next aimed to identify the characteristics of the CD4+ T cells that were sensitive to the cytotoxic action of RT53. RT53 was previously shown to induce death of cancer cells during metabolic stress (33). Cancer cells rely on high level of metabolic activities for adequate supply of biosynthetic intermediates for their life cycle, and similar conditions favor HIV-1 replication in CD4+ T-cells (7). We thus analyzed if the differential sensitivity of CD4+ T cells to RT53 treatment was related to differences in cellular metabolism. Following treatment of CD4+ T cells with RT53 but not RK16, we observed disappearance of larger CD4+ T cells (determined by FCS in flow cytometry) (Fig S6) and a decrease in the proportion of CD25^high^ and HLA-DR^high^ cells (Fig 7A). We next analyzed the metabolic activity [glycolysis, measured by extracellular acidification rate (ECAR) and oxidative phosphorylation (OXPHOS), measured by oxygen consumption rate (OCR)] of living CD4+ T cells in non-treated or cells treated with RT53/RK16 peptides (Fig 7B). We found that the cells that survived RT53 treatment had lower mitochondrial respiration (spare respiratory capacity) and glycolysis (basal ECAR and glycolytic reserve (ΔECAR after FCCP)) than cells that had not been treated or treated with RK16 (Fig 7C). We thus concluded that CD4+ T cells with the highest metabolic activity were more sensitive to the action of AAC-11-derived peptides.

**Figure 7.**
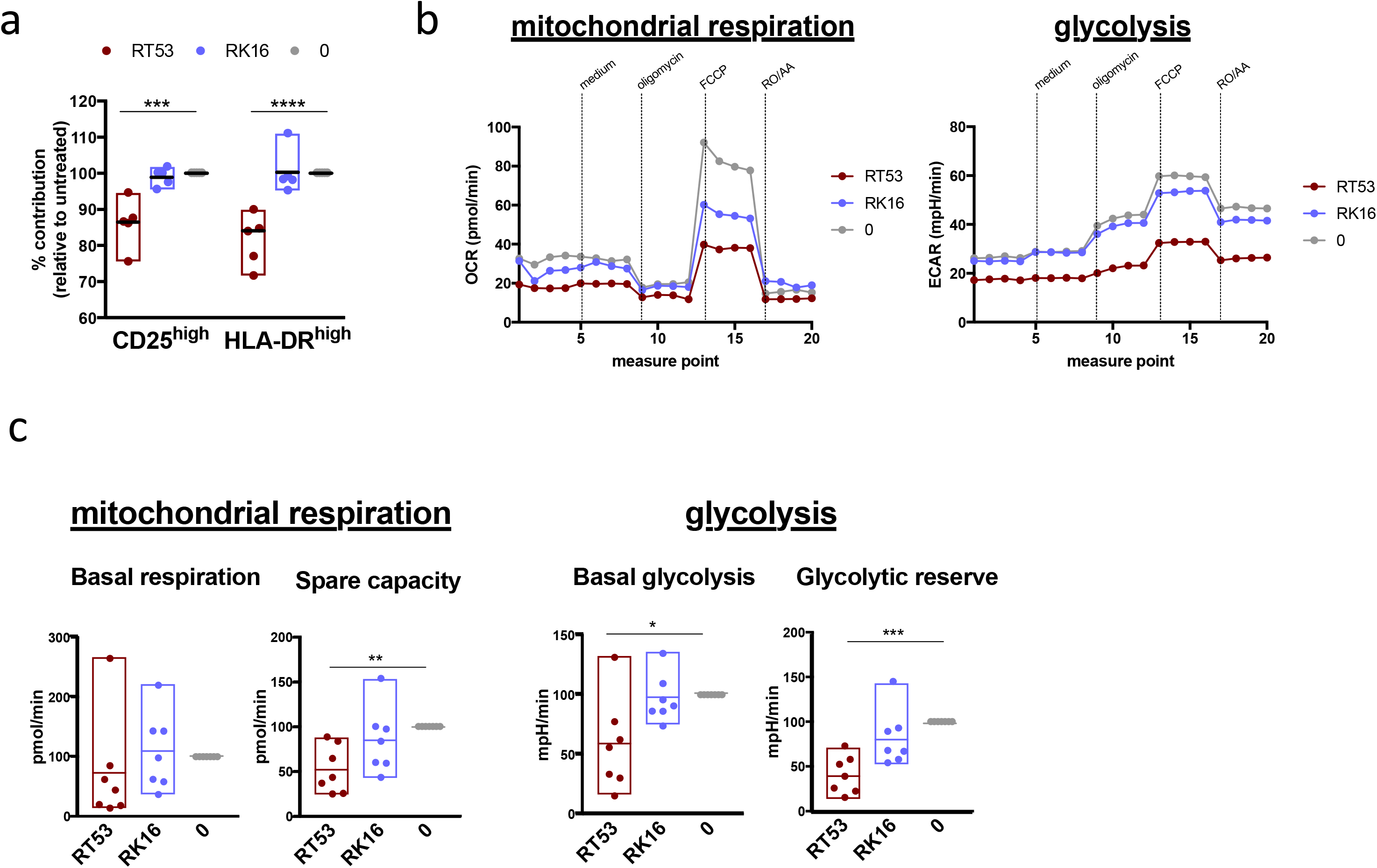
RT53 preferentially targets activated and most metabolically active cells. **A**. *In vitro* activated cells were pulsed with 6 μM RT53 or RK16 for 5 hours. The relative surface expression of CD25 and HLA-DR by flow cytometry when compared to the no peptide condition is shown. **B**. OCR and ECAR values of CD4+ T cells from one representative donor (left). **C.** Relative basal mitochondrial respiration (basal respiration), spare mitochondrial capacity (Spare capacity), basal glycolysis (basal ECAR) and glycolytic reserve (ECAR after FCCP) obtained with cells from 7 donors when compare to the no peptide condition (right). Values are expressed as min-max range. Median is indicated.

### RT53 depletes the pool of cells permissive to HIV-1 replication

To further confirm that RT53 was killing HIV-1 susceptible target cells and selecting for cells resistant to viral replication, we treated activated CD4+ T cells with RT53 or RK16 for 24 hours, sorted live cells from dead cells and infected sorted live cells (Fig 8A). We found that the cells that survived RT53 treatment were less susceptible to both HIV-1 Bal and VSVG-pseudotyped HIV-1 particles (Fig 8A), although the inhibition was more marked for the fully competent virus. Importantly, we did not see any further increase in cell death when compared to control conditions in sorted living cells.

**Figure 8.**
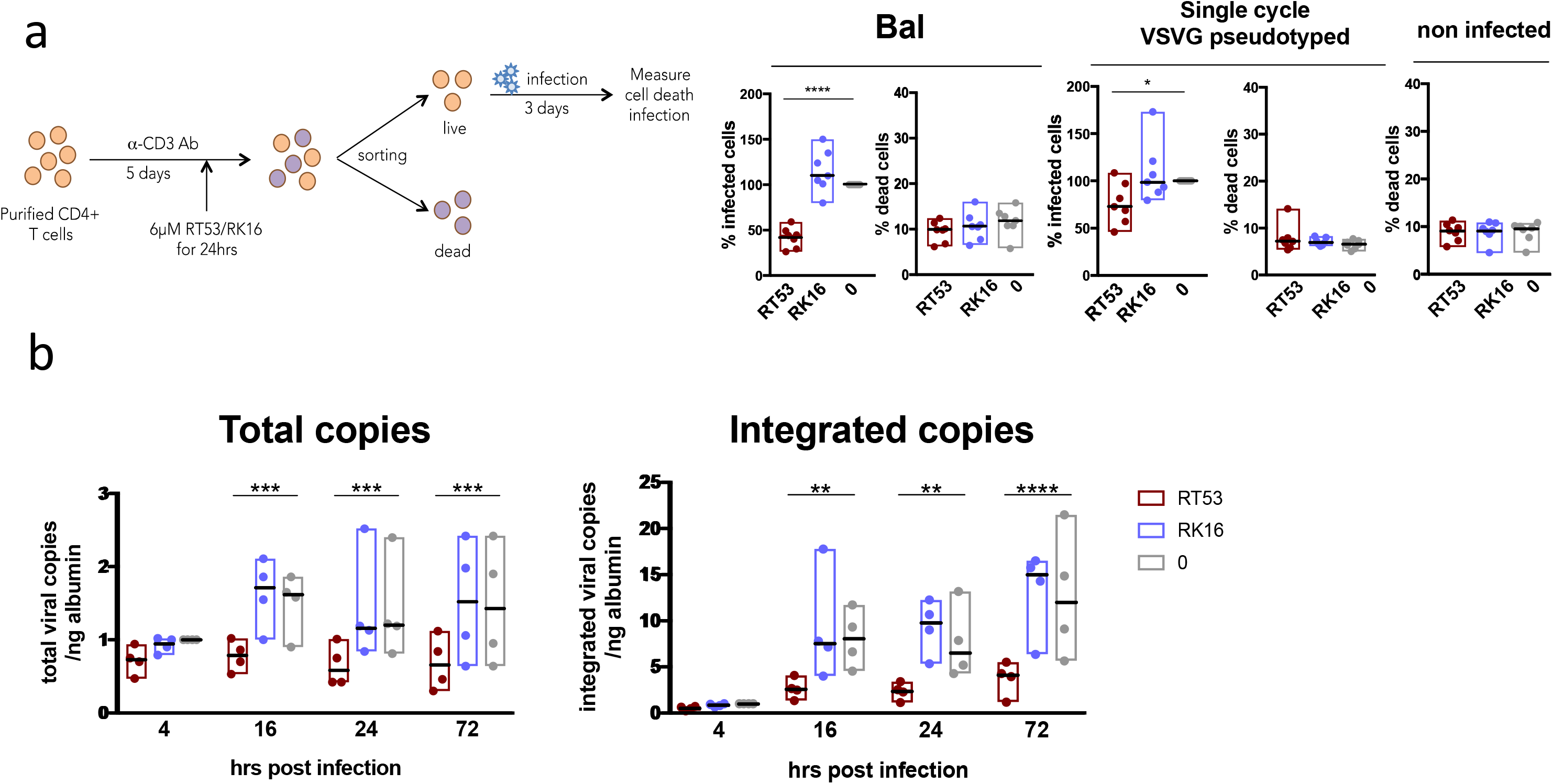
RT53-surviving cells are resistant to HIV infection. **A.** *In vitro* activated CD4+ T cells were incubated with 6 μM RT53 or RK16 for 24 hours. Live cells were then sorted from dead cells using fluorescently activated cell sorting (FACS) and live cells were infected with indicated HIV-1 strains. Cell death and infection were evaluated by flow cytometry 3 days later. **B.** Total and integrated viral copies evaluated at various time points post-infection in cells infected with VSVG-pseudotyped HIV-1 viral particles and treated with 6 μM RT53 or RK16. Values are normalized to untreated 4 hours post-infection condition and expressed as min-max range with median indicated.

We next sought to determine which stage of the viral replication cycle was blocked in cells surviving RT53 treatment. RT53 induced a decrease in the proportion of CCR5^high^ cells (Fig S7A), although this difference did not appear sufficient to explain the level of HIV inhibition found. Moreover, RT53 did not change CCR5 cell surface distribution in the cells that survived treatment with RT53 (Fig S7B). RT53 was still able to block infection when added 24h after the culture of CD4+ T-cells with HIV-1 BaL (Fig S7C). We next measured the levels of total (as a measure of reverse transcription) and integrated viral DNA at various time points post-infection with VSVG-pseudotyped HIV-1 particles in RT53/RK16 treated or untreated cells (Fig. 8B). The levels of both total and integrated HIV-1 DNA drastically increased as a function of time in RK16-treated and untreated cells but not in RT53-treated cells. These results indicate that the virus is unable to complete early steps of its life cycle in cells surviving RT53 treatment and that these cells are non-permissive to viral replication. Thus, RT53 treatment is killing cells in which the virus is able to establish productive infection, while allowing the survival of HIV-1 resistant cells.

## DISCUSSION

In this study we showed that peptides derived from the LZ region of the anti-apoptotic factor AAC-11 inhibited HIV-1 infection of primary CD4+ T cells. In particular, we found that RT53 was able to block infection with a large panel of lab adapted and primary viral strains by inducing cell death of CD4+ T cells that offered the best conditions to sustain HIV-1 replication.

Although sensitivity to RT53 was more pronounced in highly differentiated CD4+ T-cell subsets, RT53 treatment eliminated a fraction of the cells within each cell subset, rendering remaining cells resistant to HIV-1 replication. RT53 depleted highly metabolic cells, which is in agreement with the observation that a rich metabolic environment is necessary for HIV replication (7). Upon antigenic stimulation CD4+ T cells upregulate their metabolic activities, in particular glucose metabolism, to cope with the bioenergetic demands necessary to exert their functions. Thus, naïve and central memory CD4+ T-cells are typically smaller and have decreased metabolic activities when compared to more differentiated and activated subsets. However, we have recently shown that some cells with enhanced metabolic activity can be found even among phenotypically quiescent naïve T cells and these cells are susceptible to HIV-1 infection (7). Here we found that there was a close association between AAC-11 peptide-induced cell death and inhibition of HIV infection. Not only the rates of cell death induced by AAC-11 peptides correlated with the extent of HIV inhibition but precluding cell death by increasing extracellular levels of K^+^ also restored the infectability of CD4+ T cells. Cellular metabolism and cell death are deeply entangled. Metabolic arrangements are necessary to sustain long-term memory cells. External signals provided by T cell receptor activation or growth factors such as IL-2 and IL-7 promote an anti-apoptotic state of the cell by increasing the levels of metabolite transporters (e.g. Glut1) that ensure the supply of nutrients necessary to sustain the bioenergetic demands of the cell (34, 35). The absence of these signals provokes a limitation in the influx of nutrients that results in metabolic stress and the activation of cell death pathways. Our results indicate that AAC-11 derived peptides may tilt the equilibrium of highly metabolic cells towards cell death, even among less differentiated cells with a stronger anti-apoptotic basal state.

Although the anti-apoptotic action of AAC-11 is well documented, the molecular mechanisms underlying its activity are still unclear. AAC-11 possesses several domains predicted to be responsible for protein-protein interactions (19), and its expression may interfere with different mechanisms of cell death. The peptides that we tested here were designed to mimic the heptad leucine repeat region of AAC-11 and to be used as competitive inhibitors that abrogate the interaction of AAC-11 with its partners. In tumor cells physical interaction between the LZ domain of AAC-11 and Acinus prevents Acinus pro-apoptotic cleavage by caspase-3. In our study we did not observe significant changes in the expression of Acinus upon treatment with AAC-11-derived peptides (not shown) and caspase-3 activity was not significantly detected in our experimental conditions, suggesting that this pathway was not the major contributor to T cell death observed in our experiments. In contrast we found that the treatment with RT53 induced a strong increase in caspase-2 activity. Although AAC-11 has been shown to physically bind to the caspase recruitment domain of caspase-2 preventing its auto-cleavage (22), the activation of this caspase seen in our experiments is probably due to a side effect of the decrease in intracellular K^+^ levels caused by the peptides (22, 36). The inhibition of caspase-2 activity also did not lead to the restoration of cell viability, suggesting that the mechanisms of cell death induced by RT53 is independent of caspase-2 activation. RT53 was previously seen to localize at the plasma membrane compartment of cancer cells and was described to have a membranolytic mechanism of action. We have also observed the peptide’s localization to this cellular compartment in primary CD4+ T cells. The fast kinetics of the peptide’s activity also points to an action through rearrangement of pre-existing cellular factors probably found at the cell’s plasma membrane and enriched in metabolically active cells.

Previous analyses have ruled out the unspecific detergent-like cell death mechanism of RT53 (24). Indeed, we also found here that RT53 was cytotoxic to only some subsets of PBMCs thus confirming the specificity of RT53 action, possibly through a membrane partner absent from certain cells. Interestingly, we also observed that RT53 did not have anti-viral effect despite observable cytotoxicity in CD4+ T cells lines Jurkat and SupT1. Immortalized cell lines are known to have perturbed survival and metabolic pathways thus providing more favorable molecular environment for HIV-1 replication. This observation also points to the existence of a specific survival pathway used by HIV-1 in primary CD4+ T cells perturbed by RT53.

Of note, although we observed a consistent inhibition of infection with VSVG-pseudotyped HIV-1 single cycle particles upon treatment with AAC-11 peptides, the inhibition was more important when we used wild type viruses, either R5 or X4. There are several possible explanations to this. First, the additional block in HIV infection observed when we used wild type (WT) viruses could be due to a cumulative effect of multiple infection cycles. Second, cells selected upon treatment with AAC-11 may be more resistant to WT virus. For instance, some cells expressing high levels of CCR5 were depleted upon treatment with RT53. We cannot exclude either that treatment with AAC-11-derived peptides induced further viral restriction in the cells that survived. Finally, infection with WT HIV might induce cellular changes that can increase the susceptibility of infected cells to the action of AAC-11-derived peptides. For instance, HIV has been shown to induce metabolic changes in infected cells and in particular an increase in the expression of Glut1-3 (37).

Our results show that AAC-11 derived-peptides were active against CD4+ T cells that were particularly susceptible to HIV-1 infection. It remains to be established if this sensitivity to the peptides remains in persistently infected cells. However, our results suggest that the sensitivity to the action of RT53 decreased somewhat 72h after infection, suggesting HIV-1-mediated modification of the cell death pathways in infected cells, as has been shown for macrophages (38). The mechanisms by which an infected cell persists are not completely clear, as HIV has cytopathic effects that lead to the death of infected CD4+ T cells. It has been recently reported that persistent latently infected CD4+ T cells in which the virus has been reactivated are intrinsically resistant to killing by HIV-specific CD8+ T cells (39), which indicates that the persistent reservoir might be seeded in cells programmed to resist cell death. BIRC5 (a.k.a. Api4 or survivin), a member of the inhibitor of apoptosis protein family, is upregulated in CD4+ T cells during HIV-1 infection and contributes to the persistence of infected cells (40). Upregulation of BIRC5 on infected CD4+ T cells was triggered by OX40, a costimulatory receptor that promotes T cell differentiation and survival (41, 42) and contributes to the metabolic boost necessary for T cell activation (43). It is thus possible that HIV-1 exploits/activates several cellular survival programs associated with T cell activation and metabolic activity to ensure its persistence.

Although further studies will be necessary to directly evaluate this, our results suggest that HIV-1 may rely on AAC-11-dependent anti-apoptotic pathway for the establishment of productive infection in CD4+ T cells. AAC-11 gene expression, besides strongly correlating with multiple genes involved in the regulation of cell metabolism, was also correlated with the expression of RRM2 (Fig 1C), an enzyme that is critical for the de novo synthesis of dNTPs, and the depletion of which blocks HIV-1 infection in macrophages and dendritic cells (44, 45). AAC-11 mRNA levels also correlated with the expression of other genes that have been associated with the HIV replication cycle, in particular trafficking (i.e. CFL1, DYNC1H1, ACTB) and transcription (i.e. CDK9, NFKB1). This suggests that AAC-11 expressing cells may offer an ideal environment for HIV-1 replication. AAC-11 prevents apoptosis of tumor cells in conditions of nutrient deprivation and in the absence of growth factors. We can therefore speculate that the AAC-11 survival pathway might play a similar role in promoting persistence of memory CD4+ T cells and HIV infection. Like the mechanisms associated with HIV persistence, it is also unclear how some T cells survive to become long-lived memory cells. While most T cells undergo apoptosis at the end of the immune response when environmental signals wane, a few cells survive even in the absence of growth factors (46). We found that expression of the apoptosis inhibitory protein AAC-11 increased with differentiation of memory CD4+ T cells. Our results suggest that the AAC-11 survival pathway may be involved in the regulation of T cell immunity, a function that has not been previously described and deserves further detailed exploration.

Among current strategies under evaluation pursuing HIV cure, allogeneic hematopoietic stem cell transplant has been shown to drastically reduce the viral reservoir (47). However, this is a risky intervention associated with severe immune ablation. We found that AAC-11 peptides blocked HIV infection through the preferential killing of HIV-1 ‘infectable’ cells but preserved a significant fraction of T-cell immunity, and in particular the less differentiated T cells. RT53 has been shown to be efficient *in vivo* in murine models of cancer with little adverse effects. Although its clinical application in the context of HIV infection remains uncertain, our results serve as a proof of concept that selective elimination of HIV-1 targets is a possible therapeutic strategy.

## MATERIAL AND METHODS

### Peptides, antibodies and probes

Peptides (Proteogenix) were received as dry powder and reconstituted with water for use. Peptide sequences are as follows.

RT53: RQIKIWFQNRRMKWKKAKLNAEKLKDFKIRLQYFARGLQVYIRQLRLALQGKT,
RT39: RQIKIWFQNRRMKWKKLQYFARGLQVYIRQLRLALQGKT,
RL29: RQIKIWFQNRRMKWKKYFARGLQVYIRQL,
RQ26: RQIKIWFQNRRMKWKKLQYFARGLLQ,
RK16: RQIKIWFQNRRMKWKK.

Antibodies and dyes used for flow cytometry and FACS sorting are as follows: LIVE/DEAD™ Violet Viability dye (ThermoFisher), CD3-PE (clone SK7, Biolegend), CD4-A700 (clone OKT4, eBioscience), CD45RA-APC-Cy7 (clone HI100, Biolegend), CCR7-Pe-Cy7 (clone GO43H7, Biolegend), CD27-APC (clone M-T271, Myltenyi), CD25-PE-Dazzle594 (clone M-A251, Biolegend), HLA-DR-PerCP-Cy5.5 (clone G46-6, Biolegend), CCR5-PE (clone 3A9, BD), p24-FITC (clone KC57, Coulter). AAC-11 expression was analyzed by intracellular flow cytometry staining using BD Cytofix/Cytoperm™ buffer set (BD) and anti-AAC-11 primary antibody (ab65836, abcam) at 1/100 dilution followed by a secondary antibody conjugated to A674 (ThermoFisher, A21244) at a 1/1000 dilution. Caspase activity was assayed with the following probes according to the manufacturer’s instructions: caspase-1 660-YVAD-FMK probe (FLICA 660 Caspase-1 Assay Kit, ImmunoChemistry Technologies^LLC^), caspase-2 FAM-VDVAD-FMK probe (FAM-FLICA Caspase-2 Assay Kit, ImmunoChemistry Technologies^LLC^), caspase-3/7 SR-DEVD-FMK probe (SR-FLICA® Caspase-3/7 Assay Kit, ImmunoChemistry Technologies^LLC^) and caspase-8 FAM-LETD-FMK probe (FAM-FLICA® Caspase-8 Assay Kit, ImmunoChemistry Technologies^LLC^). FACS sorting was performed on BD Aria and flow cytometry acquisition on BD LSRII.

### Isolation and culture of primary human CD4+ T cells

Healthy donor blood prepared as a buffy coat was obtained from *Etablissement Français du Sang (EFS)* (agreement with Institut Pasteur C CPSL UNT, 15/EFS/023). Blood was overlayed on Ficol (EuroBio) at a ratio of 2:1 v/v blood to Ficol and centrifuged at 1,800 rpm for 30 minutes at a minimum acceleration/deceleration to obtain peripheral blood mononuclear cells (PBMCs). CD4+ T cells were then purified from PBMCs by negative selection using StemCell EasySep™ Human CD4+ T cell Isolation Kit. Cells were counted and cultured in RPMI-1640 containing Glutamax (ThermoFisher), 10% fetal bovine serum (FBS), penicillin-streptomycin (ThermoFisher) (100 U/ml) and IL-2 (Myltenyi) (100U/ml) (thereafter referred to as culture medium) at 10^6^ cells/ml in 37° degree, 5% CO_2_ humidified incubator. Cells were activated with soluble anti-CD3 (clone UCHT-1) (Biolegend) for 5 days prior to infection or analysis as previously described (27).

### Isolation and culture of primary infected macaque CD4+ T cells

Macaque splenic CD4+ T cells were obtained from Cynomologus macaques (*Macaca fascicularis)* that were imported from Mauritius island and housed at *Commissariat à l’Energie Atomique et aux Energies Alternatives (CEA)* at Fontenay-aux-Roses in France under the compliance with the Standards for Human Care and Use of Laboratory Animals (Assurance number A5826-01). These animals were part of the pVISCONTI study, which received ethics approval under the number APAFIS#2453-2015102713323361 v2. They were infected intravenously with 1000 AID_50_ of SIVmac_251_ and sacrificed at a study end point at which time spleen samples were obtained. Splenic macaque CD4+ T cells were purified by mechanical disruption of a spleen sample in RPMI media using GengleMACS™ dissociator (Milntenyi) followed by cell overlay over Ficol (EuroBio) (diluted with PBS to 90% prior to use) and centrifugation at 350g for 20min. Cells were then subjected to red cell lysis and then to CD4+ T cell negative selection using CD4+ T cell negative selection kit (Myltenyi). Cells were cultured in the culture media overnight prior to incubation with peptides. Viral spread from *in vivo* infected cells was monitored by ELISA quantification of p27 (Expressbio) levels on culture supernatants.

### Infection and peptide treatment of primary CD4+ T cells

Activated CD4+ T cells were infected with either: Bal (2.9 ng/ml p24), BX08 (21 ng/ml p24), DH12 (3 ng/ml p24), 132w (6.3 ng/ml p24), NL4.3 (7 ng/ml p24), vesicular stomatitis virus glycoprotein (VSVG) pseudotyped-Δenv-Δnef-GFP (7 ng/ml p24) or SIV_mac251_ (36.7 ng/ml p27) viruses by centrifuging at 1,200g for 1 hour at room temperature and then incubating for 1 hour at 37° degrees in a humidified 5% CO_2_ incubator. Cells were then washed once with PBS, incubated in the culture medium and treated with peptides at 6 μM concentration unless otherwise indicated. Cell death and infection were measured on day 3 post-infection unless otherwise stated. Cell death was evaluated using flow cytometry (LIVE/DEAD™ Violet Viability dye (ThermoFisher)) and infection was evaluated by either flow cytometry (intracellular p24 staining) or p24/p27 ELISA (XpressBio).

### Real-time flow cytometry

Cells were washed once with PBS and incubated in Annexin buffer (10 mM HEPES, 140 mM NaCl, 2.5 mM CaCl_2_, pH 7.4) in the presence of Annexin V-FITC (Biolegend) at a concentration of 10^6^ cells/ml for 15 minutes. 7-AAD (Biolegend) was added for the last 5 minutes of incubation. Cells were then directly passed through the flow cytometer to acquire fluorescence vs time without washing. Alternatively, cells were stained with asanate potassium green 2 (APG-2) AM (Abcam), a fluorescent K^+^ indicator, at a final concentration of 1 μM in plain RPMI-1640 medium for 30 minutes at room temperature, washed twice with plain RPMI-1640, resuspended in PBS at 10^6^ cells/ml and incubated with 7-AAD (Biolegend) for 5 minutes before acquisition. To measure mitochondrial membrane potential, cells were stained with JC-1 indicator (Thermo Fisher) at a final concentration of 2 μM in PBS at 37° C for 15 minutes, washed once and analyzed on a flow cytometer. 6 μM peptides were added after 2 minutes of baseline acquisition and acquisition continued for additional 28 minutes.

### Immunofluorescence microscopy

Activated CD4+ T cells were incubated with 6uM RT53-Rhodamine [Rhodamine-RQIKIWFQNRRMKWKKAKLNAEKLKDFKIRLQYFARGLQVYIRQLRLALQGKT] or RK16-Rhodamine [Rhodamine-RQIKIWFQNRRMKWKK] (Proteogenix) for 2 hours. Cells were then washed twice with PBS, immobilized on poly-lysine coated coverslips, fixed with 4% paraformaldehyde for 10 minutes at room temperature (RT), neutralized with 50mM NH4Cl for 10min at room temperature and washed twice with PBS. Cells were then permeabilized with 0.1% Triton-X 100 in PBS for 5 min at RT. All antibody incubations were performed in 1% bovine serum albumin (BSA) in PBS. Intracellular staining for AAC-11 was performed with anti-AAC-11 antibody (Abcam, ab65836) 1/200 dilution for 1hr at RT followed by anti-rabbit IgG-A647 secondary antibody (LifeTechnologies, A31573) 1/400 dilution for 45 min at RT. Coverslips were then washed, stained with DAPI for 15min at RT and mounted using Fluoromont G (ThermoFisher) mounting medium.

### Measurements of cellular metabolism

Oxygen consumption rate (OCR) and extracellular acidification rate (ECAR) were measured on Seahorse XF96 analyzer using Seahorse XF Cell Mito Stress Test Kit (Agilent). Briefly, activated CD4+ T cells were incubated with 6 μM RT53 or RK16 for 4 hours at 37°C. Cells were then counted, washed in a Seahorse XF medium (Agilent Seahorse XF base medium with 2mM Glutamax (Agilent, 102365-100) containing 10mM glucose (Sigma), 2mM sodium pyruvate (LifeTechnologies) and adjusted to pH 7.4. Equal number of live cells was seeded at 2*10^5^ cells per well in XF96 V3 PS plates (Seahorse Bioscience) precoated with 0.5 mg/ml Corning® Cell-Tack™ Cell and Tissue Adhesive (Corning, 354240), and incubated for a minimum of 30 minutes in a CO_2_-free, 37°C incubator prior to acquisition. The following drugs were placed at injection ports: Seahorse XF medium (port A), 2.5 μM Oligomycin (port B), 0.9 μM FCCP (port C), 1 μM rotenone and 1 μM antimycin A (port D).

### Quantitative RT-PCR

#### Evaluating the expression of AAC-11 and other genes in CD4+ T cell memory subsets

The expression levels of AAC-11 and other genes on CD4+ T cell subsets were analyzed in a previous study (7) (Dataset: DOI: 10.17632/vfj3r27gnf.1). Briefly, the total RNA from 5*10^4^ cells was extracted using RNA Trace Kit (Macherey Nagel), treated with DNase, reverse transcribed using Reverse Transcription Master Mix (Fluidigm), pre-amplified using PreAmp Master Mix (Fluidigm) and treated with exonuclease I (New England Biolabs). Samples were then mixed with SsoFast EvaGreen Supermix with Low ROX (Biorad), DNA Binding Dye (Fluidigm), and assay mix (assay loading reagent (Fluidigm) and Delta Gene primers (Fluidigm)). The expression levels were read on Biomark HQ system (Fluidigm). The expression levels of BECN1 were used for normalization. Gene expression values are plotted as 2^−ΔΔCt^.

#### Evaluating the expression of viral gene products

Cells were collected by centrifugation and dry pellet was stored at −80°C until DNA extraction. DNA was extracted using NucleoSpin® Tissue Kit (Macherey Nagel). Real time PCR (RT-PCR) to quantify total and integrated HIV-1 DNA was performed as described previously (48) using TaqMan™ Universal PCR Master Mix (ThermoFisher). Briefly, total viral DNA was quantified using the following primers and probe: CTT TCG CTT TCA AGT CCC TGT T (forward), AGA TCC CTC AGA CCC TTT TAG TCA (reverse), (FAM)-TGG AAA ATC TCT AGC AGT GGC GCC C-(BHQ1) (probe). 8E5 cell line containing a single viral copy per cell was used as a standard. Integrated viral DNA was quantified by first pre-amplifying the DNA using AccuTaq™ LA DNA Polymerase (Sigma) using the following primers: AGC CTC CCG AGT AGC TGG GA (FirstAluF); TTA CAG GCA TGA GCC ACC G (FirstAluR); CAA TAT CAT ACG CCG AGA GTG CGC GCT TCA GCA AG (NY1R). Second DNA amplification round was performed with TaqMan™ Universal PCR Master Mix (ThermoFisher) using the following primers: AAT AAA GCT TGC CTT GAG TGC TC (NY2F); CAA TAT CAT ACG CCG AGA GTG C (NY2R); FAM-AGT GTG TGC CCG TCT GTT GTG TGA CTC-TAMRA (NY2ALU probe). HeLa cells line containing HIV-1 integrated DNA was used as a standard (48). Results were normalized to ng actin using Human DNA standard (Sigma). Primers used for quantification of actin were: TGC ATG AGA AAA CGC CAG TAA (forward); ATG GTC GCC TGT TCA CCA A (reverse); (FAM)-TGA CAG AGT CAC CAA ATG CTG CAC AGA A-(TAMRA) (probe).

### Statistical analysis

Statistical analysis was performed using GraphPad Prism software. Linked parametric or non-parametric one-way ANOVA or two-way ANOVA were used depending on the experiment. * denotes p≤0.05, ** denotes p≤0.01, *** denotes p≤0.001, **** denotes p≤0.0001. Dunnett’s, Dunn’s or Holm-Sidak’s multiple comparisons tests were used for post-hoc analysis.

## Acknowledgements

The authors would like to thank all the blood donors for their generous contribution to research. Anastassia Mikhailova and Amal Elfidha were enrolled in the *école doctorale* Bio Sorbonne Paris Cité (BioSPC) for their PhD program. The authors acknowledge the Cytometry and Biomarker UTechS at Institut Pasteur and the personnel from the Infectious Disease Models and Innovative Therapies (IDMIT) plate-form for support in conducting this study.

## Financial Support

Anastassia Mikhailova was supported by the Pasteur-Paris University (PPU) International PhD Program. Amal Elfidha was supported by the ANRS. The study was supported with funds from the Institut Pasteur VALOEXPRESS program. The ANRS pVISCONTI study was supported by ANRS and MSDAVENIR.

## Author Contributions

AM, JCV-C, AD, VM and AE performed experiments; AM, JCV-C, AD, VM, SV and AS-C analyzed the data; JLP and CP provided key reagents; JLP and AS-C conceived the study; AM, AD, JCV-C and AS-C designed the experiments; AS-C supervised the study; AM and AS-C drafted the article; and all authors critically reviewed the manuscript.

